# The coming of age of EvoMPMI: evolutionary molecular plant-microbe interactions across multiple timescales

**DOI:** 10.1101/254755

**Authors:** Jessica L Upson, Erin K Zess, Aleksandra Białas, Chih-hang Wu, Sophien Kamoun

## Abstract

Plant-microbe interactions are great model systems to study co-evolutionary dynamics across multiple timescales, ranging from multimillion year macroevolution to extremely rapid evolutionary adaptations. However, mechanistic research on plant-microbe interactions has often been conducted with little consideration of the insights that can be gained from evolutionary concepts and methods. Conversely, evolutionary research has rarely integrated the diverse range of molecular mechanisms and models that continue to emerge from the molecular plant-microbe interactions field. These trends are changing. In recent years, the incipient field of evolutionary molecular plant-microbe interactions (EvoMPMI) has emerged to bridge the gap between mechanistic molecular research and evolutionary approaches. Here, we report on recent advances in EvoMPMI. In particular, we highlight new systems to study microbe interactions with early diverging land plants, and new findings from studies of adaptive evolution in pathogens and plants. By linking mechanistic and evolutionary research, EvoMPMI promises to add a new dimension to our understanding of plant-microbe interactions.

## Introduction

The field of plant-microbe interactions is remarkable for the recurrence of common processes across unrelated species and pathosystems. For example, distantly related plant pathogens exhibit extensive similarities in both their virulence and adaptation mechanisms, and conserved disease resistance toolkits and networks underlie immune responses across plant taxa. These features make plant-pathogen interactions excellent model systems for developing and challenging evolutionary biology concepts.

Plant-pathogen systems are also exceptional in that their evolutionary dynamics can be studied across multiple timescales. In macroevolutionary terms, microbes have shaped the evolution of plants since their arrival on land—and vice versa—resulting in fine-tuned interactions, specialised genome architectures, and increased robustness through redundancy and the emergence of networks. With shorter timescales in mind, the arms race that exists between plants and microbes creates a tight interplay, with the plant activating defence responses that hinder the growth of the pathogen, whilst microbes deploy specialised molecular weapons that act within the plant to provide a foothold for infection. These dynamics create strong selective pressures, and have led to striking examples of rapid adaptive evolutionary change.

Nevertheless, mechanistic research on plant-microbe interactions has often been conducted without an appreciation of the insights that can be gained from evolutionary concepts and methods. Conversely, evolutionary research has rarely integrated the rich spectrum of molecular mechanisms and models that continue to emerge in the molecular plant-microbe interactions field. This is changing. In recent years, the nascent field of evolutionary molecular plant-microbe interactions (EvoMPMI) has emerged to link mechanistic molecular research to evolutionary approaches.

In this review, we summarise recent advances in EvoMPMI. We broadly report on the trends and constraints that drive the coevolution of plants and microbes. In particular, we highlight new systems to study microbe interactions with early diverging land plants, and new findings from studies of adaptive evolution in pathogens and plants. By narrowing the gap between mechanistic and evolutionary research, EvoMPMI promises to add a new dimension to our understanding of plant-microbe interactions.

## The dawn of plant-microbe interactions

Without microbes, plants, as we know them, would not exist. Terrestrial plants are thought to have evolved as the result of an ancient symbiosis between a semi-aquatic green alga and an aquatic fungus, borne onto land over 450 millennia ago (Delaux et al. 2015). In this model, the colonization of land by plants—and therefore their very evolution—was only possible through an intimate partnership with a filamentous microorganism (Ponce de León and Montesano 2017; Carella et al. 2017; Wang et al. 2012).

Interestingly, this foundational fungal symbiosis may have come at a cost, since some microbes appear to have co-opted plant’s symbiont accommodation to establish pathogenic interactions early on during land plant evolution (Wang et al. 2012). Indeed, some of the genes required for fungal symbiosis in plants are necessary for extensive colonization by biotrophic pathogens (Wang et al. 2012; Carella et al. 2017). However, the topic appears to be somewhat controversial (Huisman et al. 2015). Nonetheless, analyses of the fossil record indicate that plant-pathogen interactions may date as far back as 400 million years ago (Mya); this consists of structures diagnostic of plant defences, such as encasements around intracellular hyphae (Krings et al. 2007). Moreover, a seed fern fossil that dates back to the Carboniferous period (300 Mya) displays a haustorium-like infection structure and is possibly the earliest evidence of parasitism by oomycetes (Strullu-Derrien et al. 2015).

Although there is evidence that plant-pathogen coevolution dates back to ∼400 Mya, the cellular and molecular mechanisms underpinning these early interactions remain obscure. One approach that has gained popularity in recent years is to study the interactions of extant microbes with early diverging land plants, notably the bryophytes—hornworts, mosses, and liverworts—and determine how these interactions differ from those with angiosperms (**Figure 1a**) (Rensing et al. 2008). The rationale is that comparative studies with a range of land plants will shed light on ancestral features of plant-microbe interactions. It turns out that many microbial pathogens can infect bryophytes, causing disease symptoms such as necrosis and tissue maceration (Ponce de León and Montesano 2017). The oomycete *Phytophthora palmivora* was shown to colonize the vascular layer of liverworts—the earliest diverging land plant—and to form intracellular hyphal structures similar to the haustoria produced on angiosperms (Carella et al. 2017) (**Figure 1b**). Liverwort cellular trafficking proteins and membrane protein syntaxins accumulate at these *P. palmivora* infection structures, suggesting that plant accommodation of oomycetes is an ancient feature of land plants (Carella et al. 2017). Other studies have examined the moss *Physcomitrella patens* response to pathogen molecular patterns and filamentous microorganisms to gain insight into the evolution of plant defences (Ponce de León and Montesano 2017; Bressendorff et al. 2016; Overdijk et al. 2016). *P. patens* perceives the presence of pathogens and activates defences that include the accumulation of reactive oxygen species, localized cell death, and induction of defence gene expression (Bressendorff et al. 2016).

**Figure 1.**
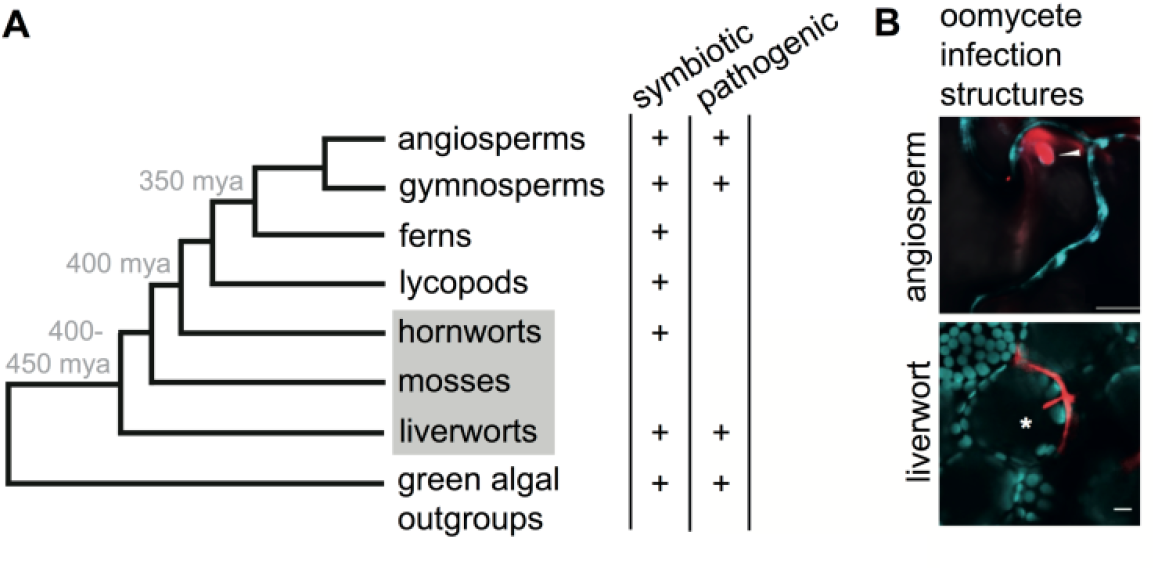
Specialized intracellular infection structures across land plants. (A) Evolution of microbial interactions in land plants. Simplified phylogeny of land plants with special attention to bryophytes (grey box) and estimated date of divergence (grey text). Evidence for specialized symbiotic or pathogenic intracellular infection structures in the various lineages is denoted by a ‘+.’ **(B) Intracellular pathogen infection structures across land plants**. Intracellular infection structures of the *Phytophthora* species *P. infestans* and *P. palmivora* on evolutionarily distant hosts, the angiosperm *Nicotiana benthamiana* (Chaparro-Garcia et al. 2011) and the liverwort *Marchantia polymorpha* (Carella et al. 2017), respectively. Each *Phytophthora* species (red) forms a haustorium (marked by an arrow or star)—a digit-like protrusion that serves as the host interface—in its respective host cell, with cell boundaries indicated by cytoplasmic markers (top, turquoise) or plastid auto-fluorescence (bottom, turquoise).

These experimental systems provide a range of exciting new platforms to study the evolutionary dynamics of plant susceptibility and immunity to microbial pathogens, and complement similar systems developed for symbiotic interactions (Ligrone et al. 2007). We can now ask pertinent questions about the macroevolutionary dynamics of plant symbiosis, pathogenesis and immunity. How did microbes evolve to infect land plants? What are the commonalities and differences in microbial accommodation by plants across symbiotic and pathogenic interactions? When and how did plant immune receptors evolve? How did immune signalling pathways develop over time? This comparative approach—combined with detailed mechanistic understanding of microbial interactions with higher plants—will shed light on the factors that have shaped these complex interactions and thus driven evolution.

## Pathogen adaptations

Plant hosts are continuously shaping pathogen evolution, driving recurrent and sustained changes within microbial genomes (Raffaele and Kamoun 2012; Croll and McDonald 2012; Dong et al. 2015). The push and pull of the natural selection forces imposed by host plants drive rapid and dynamic changes in the pathogen, ultimately shaping their genome evolution over both short and long timescales (Dong et al. 2015). Successful infection requires the pathogen to modulate plant processes, for example by suppressing plant immunity. This is achieved via the actions of pathogen-secreted virulence factors known as effectors that act directly on host molecules (Dodds and Rathjen 2010; Hogenhout et al. 2009; Win et al. 2012). Thus, pathogen adaptation to their host environment is intimately linked to the evolution of their effector repertoires. Remarkably, despite the huge diversity of plant pathogen taxa and the host species that they infect, the evolution of pathogen effectors shares many common features across diverse pathosystems.

What drives pathogen effector evolution? Two broad pressures imposed by the plant host have been defined: effectors evolve to adapt to host targets, or to evade detection by plant immune receptors (**Figure 2**). The interplay between these fluctuating selection pressures accelerates the tempo of effector evolution, and creates an inherently unsettled biotic environment for the effectors, therefore generating an evolutionarily unstable framework.

**Figure 2.**
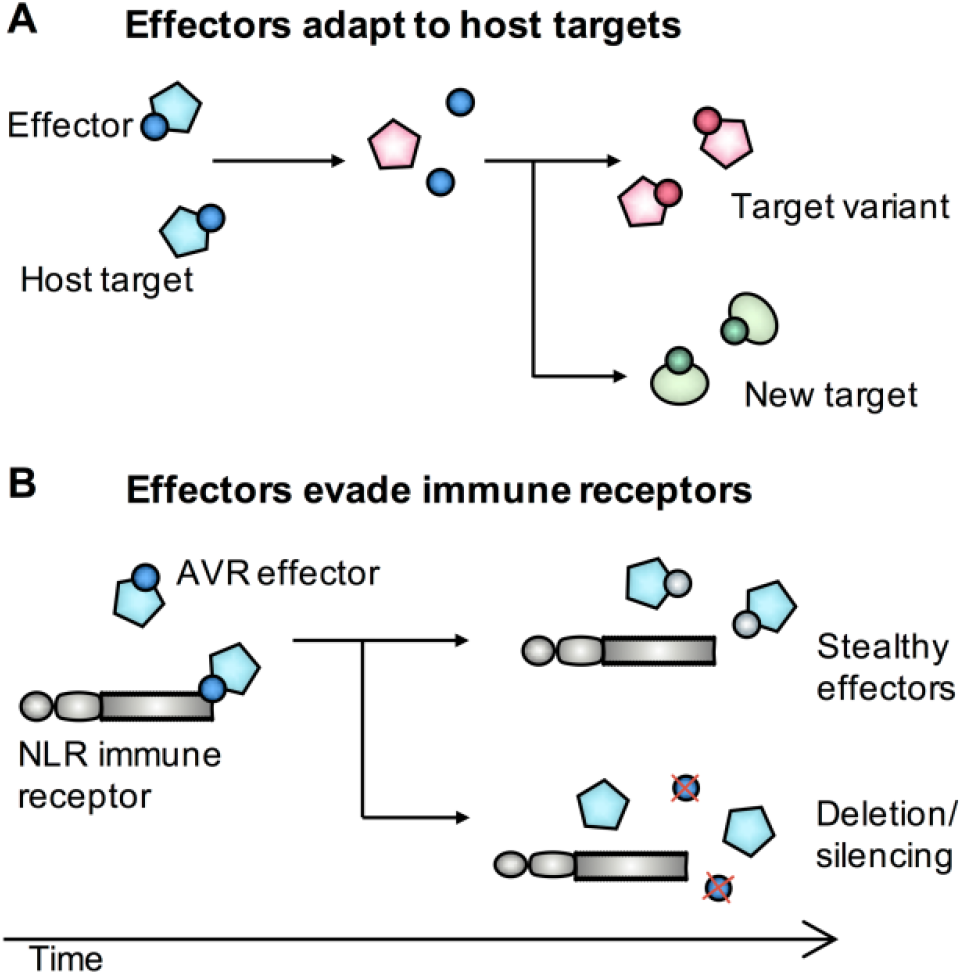
Drivers of pathogen effector evolution. **(A) Effectors adapt to new targets**. Natural variation in an effector host target or changes in the pathogen biotic environment, for example following a host jump, drive effector adaptation. This results in effectors which can bind or act on a variant of the original target or on a totally new host target. **(B) Effectors evade immune receptors**. Effectors also evolve to evade recognition by host immune receptors. This can occur through adaptive mutations that result in stealthy effectors which avoid host recognition whilst retaining virulence activity. Alternatively, effector genes can also evade host immunity through pseudogenisation, deletion, or gene silencing.

Avirulence (AVR) effectors that are detected by plant immune receptors represent remarkable examples of rapid evolutionary adaptations. Notably, stealthy effector variants that evade detection by the plant whilst retaining virulence function (**Figure 2b**) can carry extreme signatures of adaptive evolution with an excess of nonsynonymous polymorphisms (amino acid replacements) (Białas et al. 2017). Such is the case for the effector AVR-Pik of the rice blast fungus *Magnaporthe oryzae*, in which allelic variants only carry non-synonymous polymorphisms that map to the binding interface of the rice immune receptor Pik-1 (Yoshida et al. 2009; Maqbool et al. 2015). Each of the four amino acid polymorphisms of AVR-Pik appear to be adaptive (Yoshida et al. 2009; Maqbool et al. 2015), and the lack of synonymous changes is a hallmark of extremely rapid evolution, most likely driven by an accelerated arms race with the Pik-1 receptor (Białas et al. 2017). In addition, effectors also exhibit marked patterns of selection following host jumps, where there is extreme pressure to adapt to new host targets (**Figure 2a**). For example, orthologous protease inhibitor effectors from the Solanaceae-infecting *Phytophthora infestans*, and its sister species, *Phytophthora mirabilis*, which infects *Mirabilis jalapa*, have adapted and specialized to protease targets unique to their respective host plants (Dong et al. 2014) (**Fig. 3a**).

**Figure 3.**
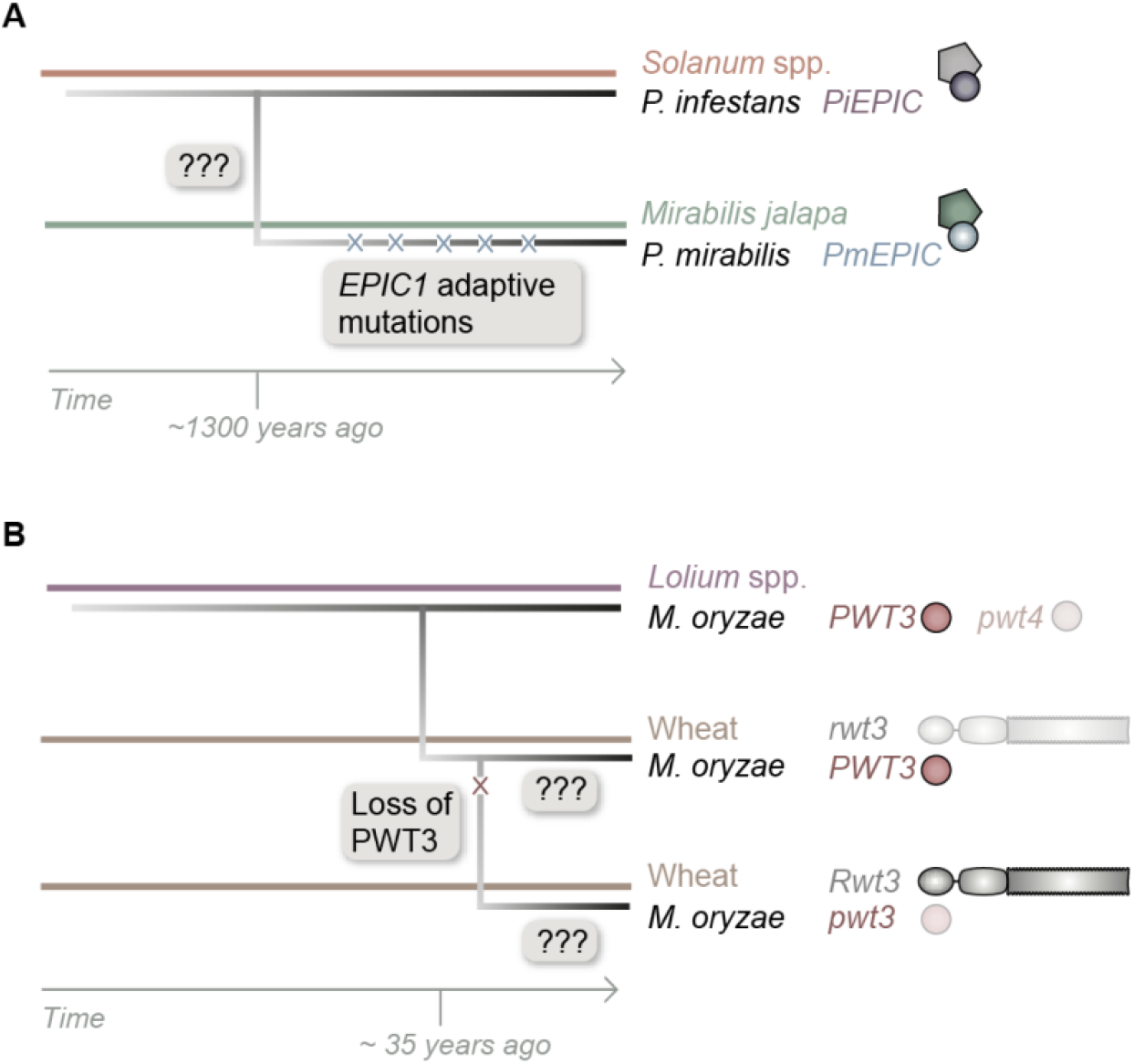
Pathogen mutations associated with host jumps. **(A) Adaptive mutations that accumulate after a host jump**. Pathogens acquire mutations that fine-tune the pathogens interaction with its new host, such as in the case of the adaptive mutations in the *Phytophthora mirabilis* protease inhibitor effector *PmEPIC*, which arose to target a *Mirabilis jalapa* host protease (Dong et al. 2014). Ultimately, such mutations lead to specialisation through inadvertent accumulation of mutations that are maladaptive in the ancestral host, in this case the *Solanaceae*. In this example, due to the relative distance of the evolutionary event, the causal mutations that enabled the host jump cannot be elucidated. **(B) Mutations that enable a host jump**. Serial loss of the *Magnaporthe oryzae* effectors PWT4 and PWT3 enabled the evolution of wheat-infecting lineages that are not constrained by the major determinants of wheat resistance, *Rwt4* and *Rwt3* (Inoue et al. 2017). Although these mutations are known to have occurred ∼35 years ago, the adaptive mutations that have subsequently accumulated in these lineages to fine-tune *M. oryzae-*wheat interaction are not yet known.

Effectors also evolve through a birth and death process, resulting in significant levels of presence/absence polymorphisms within pathogen populations (**Figure 2b**). The maintenance of complex and possibly redundant effector repertoires could prove beneficial for a pathogen population, given that AVR effectors constrain the pathogen host range and their loss could enable the pathogen to jump to new hosts. A recent study on the emergence of wheat blast in Brazil in the 1980s vividly illustrates this model, and highlights the importance of studying effector evolution (Inoue et al. 2017). Losses of the effectors PWT3 and PWT4 in *M. oryzae* strains that infect the grass species *Lolium* were sufficient to enable a jump to wheat, which carries the matching resistance genes *Rwt3* and *Rwt4* (Inoue et al. 2017). The effector losses likely occurred in a stepwise manner, as *PWT4* is absent or non-functional in most of the wheat-infecting isolates, whereas only a subset of these isolates also lack *PWT3*. The authors proposed that the widespread cultivation of wheat cultivars that lack *Rwt3* in Brazil prior to the emergence of wheat blast served as a stepping stone for the pathogen to become fully virulent on the majority of wheat cultivars through stepwise mutations in the effectors (**Fig. 3b**). Following the host jump, wheat blast emerged as a destructive and pandemic disease of wheat, having spread from South America to South Asia where it’s viewed as a serious threat to food security (Islam et al. 2016).

In addition to changes in individual effector genes, global patterns of pathogen genome evolution have been noted and may be linked to macroevolutionary trends, such as a high potential for host jumps (Raffaele and Kamoun 2012; Dong et al. 2015). Independent lineages of biotrophic plant pathogenic fungi and oomycetes have highly expanded, repeat-bloated genomes (Haas et al. 2009; Rouxel et al. 2011; Raffaele and Kamoun 2012). Unrelated pathogens, such as the oomycete *P. infestans* and the fungus *Leptosphaeria maculans*, have converged towards an unusual genome architecture, in which rapidly evolving genes, particularly effectors, are localized to repeat-rich, gene-sparse compartments that contrast with the repeat-poor, gene-dense parts of the genome that carry housekeeping and more conserved genes (Dong et al. 2015; Soyer et al. 2014; Haas et al. 2009; Raffaele et al. 2010). This “two-speed genome” architecture was proposed to accelerate adaptive effector evolution through multiple mechanisms (Raffaele and Kamoun 2012; Dong et al. 2015; Soyer et al. 2014). For example, proximity of effector genes to transposable elements enhances the rate of evolution of these genes, and may even increase the propensity of these genes to get silenced—an alternative mechanism to evade host immunity (**Figure 2b**) (Na and Gijzen 2016). Also, the bipartite genome architecture may facilitate coordinated upregulation of effector genes during infection, possibly through epigenetic modifications (Raffaele and Kamoun 2012; Soyer et al. 2014). Indeed, an important research goal is to determine the mechanisms that underpin epigenetic variation in rapidly evolving “two-speed” plant pathogen genomes. Recently, Chen et al. showed that 6mA methylation in *Phytophthora* genomes is more prevalent in gene-sparse regions of the genomes, implicating this epigenetic mark in adaptive evolution (Chen et al. 2017). However, future work is needed to determine the degree to which DNA methylation and other epigenetic modifications contribute to global patterns of pathogen genome evolution.

The patterns of genome and effector evolution discussed above may not just apply to pathogenic microbes, but could also be relevant in plant-infecting insects. A recent analysis of the genome and transcriptome of the broad host range green peach aphid *Myzus persicae* revealed that clonal individuals can colonise distantly related hosts through differential regulation of sets of clustered genes (Mathers et al. 2017). The mechanisms underlying this rapid transcriptional plasticity remain unknown, but could involve some form of epigenetic modification (Mathers et al. 2017). It will be particularly interesting to compare insect strategies for infection to those of microbial plant pathogens. New insights into the evolution of the Mp10 effector of the green peach aphid points to marked differences with microbial pathogen effectors. Whereas the majority of plant pathogen effectors have rapidly diversified with little sequence and functional conservation (Franceschetti et al. 2017), Mp10 is relatively conserved across hemipteran insects and belongs to the widespread chemosensory protein 4 clade (CSP4) (Drurey et al. 2017). Remarkably, the immunosuppression activity of Mp10 evolved via gain-of-function mutations over 250 million years ago, prior to the divergence of plant-sucking insect species (Drurey et al. 2017). As more effectors are discovered in aphids and other plant parasitic insects, it will be interesting to determine whether they would exhibit similar evolutionary trends.

## Host adaptations

In the coevolutionary waltz between pathogens and plants, pathogens also shape the evolution of their hosts, leaving marked footprints on plant genomes (**Box 1**). The highly adaptable effector repertoires of plant pathogens ultimately shape the plant immune system, leading in most cases to an extremely diverse immune receptor repertoire that confers robust disease resistance against the majority of plant pathogens. This is particularly evident for the nucleotide binding leucine-rich repeat (NLR) class of proteins—the intracellular immune receptors that detect pathogens and trigger an effective immune response (Jones et al. 2016). NLRs are present in all land plants, including early diverging species (Xue et al. 2012). Hundreds of millions of years of plant-pathogen coevolution are apparent in NLR gene repertoires, which vary considerably in number and structure. With the availability of high-quality plant genomes and advances in functional characterization of plant immunity, studies on NLR evolution and adaptation have re-emerged as active research topics.

The role of effectors in shaping the evolution of immune receptors is best illustrated by the recent discovery that non-canonical domains of NLRs originate from host proteins that are targeted by effectors. Although the majority of NLRs have a conserved modular architecture, a subset of them carry these uncoventional integrated domains (IDs) (Kroj et al. 2016; Sarris et al. 2016). Examples of NLR-ID fusions include rice Pik-1 and RGA5 proteins, which carry integrated HMA (RATX1) domains (Cesari et al. 2013; Maqbool et al. 2015), rice Pii-2 with a NOI/RIN4 domain (Fujisaki et al. 2017), and *Arabidopsis* RRS1 which carries a WRKY domain (Sarris et al. 2015). IDs are thought to be derived from effector-associated host proteins, which then act as baits for effector recognition within NLRs (Cesari et al. 2014; Ellis 2016; Wu et al. 2015; Fujisaki et al. 2017). NLR-ID fusions may have provided an evolutionary advantage by detecting multiple effectors that converge on the same host target. Nevertheless, the number and diversity of IDs that have been reported in plant NLRs is astounding, and points to a highly successful evolutionary strategy of receptor diversification (Kroj et al. 2016; Sarris et al. 2016). The mechanisms underpinning the emergence of NLR-ID fusions are poorly understood. Bailey et al. recently proposed that ectopic recombination is a major driver of domain integration in *Poaceae* (Bailey et al. 2017). A “major integration clade”, which includes *RGA5, Pi-ta* and *Rpg5*, underwent repeated independent integration events (Bailey et al. 2017). This suggests that some NLR loci may be more prone to recombination that favours integration, and thus serve as hotspots for the emergence of novel NLR-IDs. More work is needed to determine the frequency and mechanisms that drive the evolution of NLR fusions, as well as the degree to which novel fusions are maintained in plant populations.

NLR expansion and neofunctionalisation is another mechanism that enables plants to keep up with pathogens. NLR genes are often located in gene clusters that can act as a reservoir of genetic diversity (Jupe et al. 2012; Zhou et al. 2004; Meyers et al. 2005). Examples of NLR genes that have duplicated include members of the highly divergent *Mla* powdery mildew fungus resistance locus in barley (Wei et al. 2002). NLR gene expansion is driven by ectopic recombination, unequal crossing over and transposition (Innes et al. 2008; Kuang 2004; Zhou et al. 2004). For instance, a recent comparative genomics analysis revealed that retrotransposition may have facilitated NLR expansion in *Solanaceae* (Kim et al. 2017). Among the NLRs they examined, tomato *I2*, potato *R3a* and pepper *L* genes appear to have originated from a single gene by retrotransposition followed by divergent evolution in each plant lineage to respond to different pathogens (Kim et al. 2017). Interestingly, NLR genes are often in close proximity to various types of transposons, and the number of NLR gene clusters positively correlates with transposable element density (Li et al. 2010), raising the possibility that transposon-driven duplications may be common in plants (Wei et al. 2002; Wicker et al. 2010). It is fascinating that NLR gene distribution and linkage to repetitive elements appears to mirror patterns of effector gene localization in plant pathogen genomes (see above).

Another emerging paradigm is that NLRs can function cooperatively to form genetic networks of varying complexity. NLRs pairs are functionally specialized, with a sensor NLR that perceives a pathogen effector requiring a helper NLR to activate immune signalling (Bonardi et al. 2011). Interestingly, each of the NLR-IDs that have been characterized to date are genetically and functionally linked to a helper NLR (Białas et al. 2017; Cesari et al. 2014). It is possible that paired NLRs have an increased tolerance to integration of non-canonical domains given that the ID is less likely to perturb the function of an NLR pair than of a singleton. NLR proteins can also form sophisticated signalling networks. A clear example is the NRC network, which has greatly expanded in *Solanaceae* and related species (**Box 1**), and exhibits an architecture where sensor NLRs rely on a few genetically downstream NRCs (<u>N</u>LR <u>r</u>equired for <u>c</u>ell death) to establish disease resistance (Wu et al. 2017). The NRCs and their sensor mates are phylogenetically related and have most likely evolved from an NLR pair that dramatically expanded in asterid plants about 100 Mya. Interestingly, the sensor NLRs underwent a much more significant expansion compared to the NRCs, which have diversified at a slower pace possibly due to constraints imposed by downstream signalling components. Convergence of immune signalling on a few NRCs may have facilitated the capacity of sensor NLRs to neofunctionalise and keep up with rapidly evolving pathogens. In this model, the classic bow-tie architecture of the NRC signalling network may enable higher evolutionary plasticity, and is reminiscent of the networks of plant cell-surface immune receptors and animal Toll-like receptors (Oda and Kitano 2006; Smakowska-Luzan et al. 2018).

### Box 1. Distinct evolutionary trends of NLR genes in mammals vs. plants.

Although NLR genes are present both in mammalian and plant genomes, the evolutionary trajectories of their repertoires have been strikingly different. First, mammals have smaller NLR repertoires compared to plants. For instance, the human and mice genomes carry 22 and 30 NLR genes, respectively, whereas in plants that number can range up to 402 in wheat (Baggs et al. 2017; Jones et al. 2016). Second, patterns of NLR evolution are different between mammals and plants across similar evolutionary timescales. For example, the divergence time for humans and mice is estimated at around 80-100 Mya, similar to the divergence time between tomato and coffee. Nonetheless, despite this relatively long timescale, human (illustrated in yellow) and mouse (grey) NLRs are typically conserved with clear orthologous relationships (Hibino et al. 2006; Stein et al. 2007). In sharp contrast, there is usually little orthology among NLR genes across plant taxa such as tomato and coffee. For example, the NRC network of NLR genes has dramatically expanded in asterid plants from an NLR pair, with at least 47 members in tomato (red) and nearly 80 in coffee (green) scattered across the genomes of these species (Wu et al. 2017). Accelerated evolution of the NRC network reflects massive functional diversification as the expanded genes confer resistance to pathogens and pests as divergent as viruses, bacteria, oomycetes, nematodes, aphids and whiteflies.

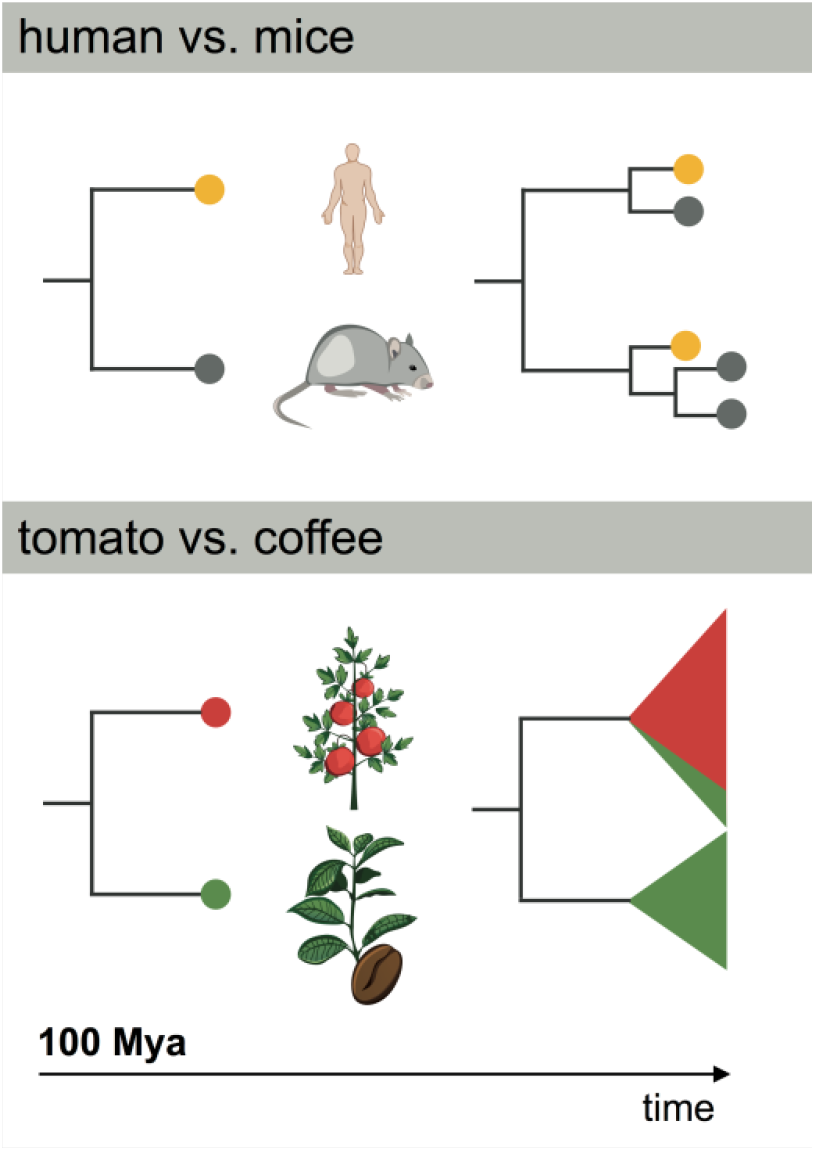

Why have mammalian and plant NLRs experienced strikingly different evolutionary trajectories across similar timescales? The answer may lie in the observation that NLRs from the plant and animal kingdoms tend to detect different types of pathogen molecules. Whereas mammalian NLRs typically detect fairly conserved pathogen molecular patterns (i.e. microbe-associated molecular patterns or MAMPs) (Stein et al. 2007), plant NLRs have evolved to detect pathogen and pest effectors that are delivered inside the host cell and are among the most rapidly evolving genes in microbial genomes. Thus, the evolutionary potential of the pathogens and their effectors drive the diversification of NLR genes, ultimately shaping the evolution of the host genome as noted with the dramatic expansions of NLR genes in plants (Wu et al. 2017).

Is NLR diversity only driven by antagonistic coevolution with pathogens? There is good evidence that NLR diversification may lead to autoimmunity and deleterious effects on plant fitness, thus restraining their evolutionary dynamics. NLR autoimmunity could be due to incompatible intramolecular interactions or interactions with other host proteins, including other NLRs. For example, Arabidopsis NLR proteins DM1 and DM2d trigger spontaneous cell death and hybrid necrosis when expressed in the same genetic background (Bomblies et al. 2007; Tran et al. 2017; Chae et al. 2014). This indicates that the evolution of the plant immune system is not only shaped by pathogen selective pressures, but also by intrinsic constraints imposed by the genetic context. It is remarkable that NLR genetic incompatibility has been observed within populations of the same species, which highlights the extremely dynamic nature of NLR evolution. The high rate of pseudogenisation observed in NLR genes further points to potential fitness penalties associated with disease resistance (Li et al. 2010; Marone et al. 2013). In addition, overexpression of some NLRs can result in autoimmunity (Heidrich et al. 2013), and NLR gene expression is generally thought to be tightly regulated through chromatin modifications and transcription factors (Zhang et al. 2016; Lai and Eulgem 2017). Furthermore, miRNAs may have also evolved to maintain homeostasis of expanded genes by targeting sequence-related NLR genes (Zhang et al. 2016). In summary, the evolutionary dynamics of plant NLRs are complex and are modulated by fitness trade-offs.

## Outlook: the coming of age of EvoMPMI

We hope that the concepts and approaches discussed above will inspire further investigations that bridge the gap between evolutionary and mechanistic molecular research of plant-microbe interactions. Performing mechanistic research with an evolutionary perspective pushes the field of plant-microbe interactions beyond the molecular, laying the foundation to ask questions not only about how these systems function, but also about how they came to be that way. Many experimental systems would greatly benefit from comparative approaches performed within a phylogenetically and ecologically robust framework to test specific hypotheses about how evolution has tweaked mechanisms of pathogenicity, symbiosis, and immunity. This approach is especially robust whenever signals of adaptive evolution are detected. Such cases complement classical experimental approaches because they imply that natural selection has performed a genetic screen in the wild under relevant biological and ecological conditions. Thus, evolutionary approaches test mechanistic molecular models under real world scenarios bringing a certain degree of rigor to our understanding of plant-microbe systems.

## Acknowledgements

We are thankful to past and present members of the Kamoun Lab as well as to several colleagues for numerous discussions and ideas. We also thank Ian Malcolm for his inspiration, and Russell Vance for a discussion that inspired Box 1. AB receives funding from the Norwich Research Park, Doctoral Training Partnership (BBSRC, UK). JU receives funding from the Gatsby Charitable Foundation. Our laboratory is funded by the Gatsby Charitable Foundation, Biotechnology and Biological Sciences Research Council (BBSRC, UK), and European Research Council (ERC- NGRB and BLASTOFF).

